# Leaf hydration status under drought is predominantly linked to stomatal regulation and leaf roll but not osmotic adjustment in Canadian hard red spring wheat (*Triticum aestivum*) cultivars

**DOI:** 10.1101/2023.03.05.531113

**Authors:** G Sharma, GS Brar, T Knipfer

## Abstract

For wheat (*Triticum aestivum*), sustained crop yield at limited soil water has been linked to osmotic adjustment (OA) as one of the main drivers to minimize drought-induced reductions in leaf hydration status and growth. Canada Western Red Spring (CWRS) cultivars are typically grown in rainfed areas in northern regions with milder climates, but ongoing climate change has increased the frequency and intensity of drought event questioning how successful they are in tolerating drought. The extent of OA and its relation to stomatal behavior, leaf roll, and kernel development under periods of drought remain elusive for CWRS. For several commercially used Canada Western Red Spring (CWRS) wheat cultivars (‘Superb’, ‘Stettler’, ‘AAC Viewfield’), OA was not found to be a mechanism contributing to drought tolerance. In contrast, we found that sustained kernel weight during periods of relatively low soil water content was linked to ‘tight’ stomatal behavior (i.e., efficient transition from onset to full stomatal closure) and ‘early’ leaf roll (i.e., reductions in flag leaf width). Moreover, leaf hydration status (Θ_RWC_) marked the onset of drought-induced losses in kernel weight in all three cultivars. In conclusion, CWRS wheat lacks OA but leaf stomatal behavior and leaf rolling aid in securing leaf hydration status and kernel weight under drought.

**One-sentence summary:** Select wheat cultivars maintain leaf hydration status and yield by early leaf roll and rapid stomatal closure in the absence of osmotic adjustment and isohydric behavior.

## INTRODUCTION

Limitations in soil water availability by drought has become the most important constraint to agriculture globally with devastating impacts on wheat production (Henson et al., 1989; Xoconostle-Cazares et al., 2011). In relatively northern regions of the world with mild summers, CWRS is commonly grown in rainfed cropping systems (De Jong et al., 2008). However, ongoing climate change has resulted in unprecedented drought events in those regions resulting in suboptimal soil water availability during the growing season which makes it a restricting factor for growing wheat with satisfactory productivity (Campbell et al., 2007). For wheat, the ability to maintain grain yield at limited soil water availability (i.e., drought tolerance) has been found to be linked to OA (Morgan and Codon, 1986; Fisher et al., 2005; Blum, 2017; Mahmood et al., 2020). Osmotic adjustment minimizes cell water loss through active solute accumulation and a decline in xylem pressure is considered as the trigger for this process (Turner and Jones, 1980; Kramer and Boyer, 1995; Turner, 2017). A major benefit of OA is maintenance of growth under drought by Ψ_P_ maintenance (Kramer and Boyer, 1995). Solute concentration effects by cell dehydration need to be distinguished from ‘active’ solute accumulation to obtain reliable estimates of OA (Babu et al., 1999, Bagatta et al. 2008). Differences in OA among wheat cultivars have been described (Morgan, 1977; Blum et al., 1999), and several wheat cultivars (such as ‘Capelle Desprez’ and ‘Condor’) only exhibit partial or no OA in response to a decline in soil water availability (Morgan, 1980; Turner and Jones, 1980; Blum et al., 1990). This raises the question: is drought-tolerance of CWRS cultivars linked to OA?

Leaf stomatal regulation plays a key role in securing plant hydration status in response to a decline in soil water availability. A drought-induced reduction in stomatal conductance reduces water loss by transpiration through stomata, which can minimize the buildup of negative leaf xylem pressures (i.e., leaf Ψ) under drought (Kramer and Boyer, 1995; Tardieu et al., 2018). Plants can be categorized into ‘isohydric’ plants that are able to maintain a nearly constant leaf Ψ until showing signs of severe dehydration and ‘anisohydric’ plants that exhibit significant reductions in leaf Ψ in repose to a decline in soil water availability (Tardieu and Simoneau, 1998). According to Henson et al. (1989), wheat cultivars (‘Gamenya’ and ‘Warigal’) exhibit anisohydric behavior because they are not able to maintain leaf Ψ and stomatal conductance under a decline of soil water content. Therefore, the question can be asked: do those CWRS cultivars that lack OA have the ability to maintain leaf Ψ under drought (i.e., isohydric behavior) through effective stomatal behavior under drought? Although other definitions of isohydricity have been used in the past, such as describing early or late stomatal closure under increasing drought stress, in this study we will refer to isohydricity in the context of leaf Ψ maintenance (Hochberg et al., 2018).

Moreover, leaf rolling can have positive impacts in minimizing water loss from the leaf surface. For example, leaf rolling reduces transpiration through the creation of a microclimate of higher humidity allowing stomata to remain in an open state to maintain CO_2_ assimilation (Oppenheimer, 1960; Kadioglu and Terzi, 2007). The microclimate is produced when moisture is trapped, and the air is cooler in the rolled leaf creating a higher relative humidity and thereby allowing the stomata to remain open (Jones, 2008). For CWRS wheat cultivars, Willick et al. (2018) reported that improved drought tolerance of cultivar ‘Stettler’ compared to ‘Superb’ is associated with greater leaf rolling and leaf relative water content. However, the authors concluded on differences in leaf rolling between cultivars following excision of leaves from the plant, which excludes any possible conclusions on the coordination of leaf rolling, stomatal regulation, and OA in the intact plant.

For commercially used CWRS wheat cultivars ‘Superb’, ‘Stettler’ and ‘AAC Viewfield’, the goal of this study was to elucidate the impact of water stress by drought (i.e., reduction in soil water content) on leaf water relations (i.e., relative water content, water potential, solute potential, stomatal conductance, and rolling) and thousand kernel weight. The data shown here highlight cultivar-specific differences in drought responses depending on the level of soil water content plants experiencing during drought.

## MATERIAL AND METHODS

### Plant material and growth

Three Canadian CWRS wheat cultivars ‘Superb’ (Townley-Smith et al., 2010), ‘Stettler’ (DePauw et al., 2009) and ‘AAC Viewfield’ (Cuthbert et al., 2018) were grown in large 10.6-L cylindrical pots. Pots were constructed from PVC tubes (15 cm x 60 cm; diameter x height, respectively) and a metal mesh (pore size around 4 mm^2^) was secured to the bottom end with a hose clamp to retain the soil medium. Each pot was equally filled with soil mix (two parts industrial sand with particle size ranging from 0.2 to 0.6 mm and one part greenhouse potting mix consisting of 75% peat and 25% perlite) up to a height of around 56 cm. This type of pot size and soil medium combination was chosen to minimize spatial inhibitions to root system growth and facilitate drainage; pots were placed on the ground and raised on plastic platforms (around 4 cm in height) to aid drainage following watering. For each pot, two seeds per cultivar were placed about 2 cm below the topsoil layer to initiate germination. One week after germination, one seedling was left in each pot and the other was pulled by hand. Two weeks after germination, about 20 g of slow-release fertilizer (Osmocote 14-14-14) was added to each pot to provide a mix of macro-/micronutrients facilitating plant growth. From time of seeding until early anthesis, plants were maintained well-watered by watering with tap water every 2-3 days to about 90% soil relative water content (soil RWC; for details see below) until the early anthesis stage (Zadoks 61, heads started to gain florets at around 60 days after seeding). When a majority of the heads had started flowering, the plant was considered to be in early anthesis. Consequently, when over 90% of the plants reached early anthesis, drought experiments were initiated. Any plants which had not headed were removed from the experiment.

Two consecutive drought experiments were performed, i.e., drought experiment 1 (duration of 19 days, seeding in early June 2021, initiation of experiment in early August 2021) and experiment 2 (duration of 17 days, seeding in mid-February, initiation of experiment in late April 2022) (see Supplementary Figure S1 for environmental conditions and Supplementary Figure S2 for plot design). During drought experiment 1, daily minimum and maximum temperature ranged from 18°C to 28°C, relative humidity from 50% to 90%, and vapor pressure deficit (VPD) from 0.30kPa to 1.50kPa (Supplementary Figure S1); during drought experiment 2, temperature ranged from 18°C to 23°C, relative humidity from 35% to 65% (relative humidity), and VPD from 0.75kPa to 1.65kPa during the drydown (Supplementary Figure S1). For both drought experiments plants were arranged following a randomized block design (see Supplementary Figure S2). In drought experiment 1, control plants were well-watered (n=6 per cultivar) and drought plants were subjected to a progressive drydown by not watering (n=6 per cultivar, Supplementary Figure S3 for soil RWC values over the drydown). The main purpose of drought experiment 1 was to determine leaf water status and associated threshold Θ_RWC_ (=onset of severe leaf dehydration) and the ability of plants for leaf osmotic adjustment in response to a progressive decline in soil RWC. Parameters measured (daily) during experiment 1 were soil RWC, leaf RWC, leaf Ψ, and leaf Ψ_S_. Following experiment 1, and using information on Θ_RWC_, for drought experiment 2 plants (n=3 plants per cultivar and watering treatment) were maintained for 17 days at 90%, 60% and 45% soil RWC (Supplementary Figure S4 for soil RWC values over the drydown); the main purpose was to determine the effect of a limited soil water availability on factors that determine leaf water loss and kernel weight. Parameters measured (every 1-3 days) during experiment 2 were soil RWC, Gs, FLW, and TKW. Considering daily time constraints when collecting data and logistical reasons, it was decided to rather measure a selected suite of parameters for each experiment and inform experiment 2 from observations in experiment 1 (for details on measured parameters see below).

### Soil relative water content

Soil RWC was determined from measurement of total pot weight (Eq. 1; Knipfer et al. 2020b):

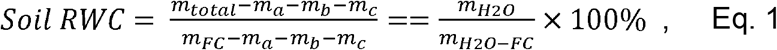

For each plant, m_total_ was measured daily (between 12 to 2 pm) using a digital balance (Uline Industrial Platform Scale) and following measurements of leaf water relations (as described below). For determination of m_FC_ (total pot weight at field capacity), the pot was fully saturated at day-0 of the drought experiment, allowed to drain for 1-2h, and measured at field capacity. Parameters of m_a_ (plant weight, on average 100g of n=10 plants), m_b_ (weight of plastic pot, on average 2.05kg), and m_c_ (soil dry weight) were treated as constants. Increases in plant weight of the experimental period were less than 750g which only contributed to a change in soil RWC of less than 7%. Soil dry weight was calculated from m_H2O-FC_ (water weight at field capacity) and soil water holding capacity at field capacity (WC_FC_ = 0.36); WC_FC_ was measured by weighing (Ohaus TP600S) soil samples at field capacity of approximately 500g in size (m_FC’_), drying soil samples in the oven at 70°C for 24-h, weighing again (m_dry’_), and calculated according to (m_FC’_ – m_dry’_) / m_dry’_) (i.e., soil gravimetric water content at field capacity). In addition to soil RWC, soil gravimetric water content (soil GWC) was determined from soil core samples at 3 different depths (15, 30, 45 cm from top) of the pot during experiment 1 (Supplementary Figure S5) to investigate soil water distribution along the length of the pot at a given soil RWC (Appendices 6 and 7). Soil core samples were extracted horizontally using a metal straw and placed in a 20 mL sealed scintillation vial; soil cores ranged from 20g to 25g and were stored in a cooler until they were transported to lab and measured within 2-h after sampling.

### Leaf water potential

Leaf water potential was measured on a flag or penultimate leaf (see Supplementary Figure S6 for measurement procedure). Prior excision from the leaf base, the leaf was bagged (>30min) in a sleeve made from aluminum foil to equilibrate leaf apoplast with symplast; measurements of an unbagged leaf results in errors caused by ongoing transpiration and disequilibrium conditions (Turner, 1981). Immediately after excision, the leaf was sealed in a second plastic bag to maintain equilibrium conditions, stored in a cooler filled with ice (i.e., avoid evaporation from leaf), and transported to the laboratory. Within 1-h of excision, leaf Ψ was measured using a Scholander-type pressure chamber (PMS Model 615 Pressure Chamber Instrument). For measurement of leaf Ψ, the top portion of the leaf was cut (length of 6 to 10 cm) and inserted through the chamber lid (i.e., bottom portion was used for leaf RWC and solute potential measurements as described below). The pressure in the chamber was raised slowly and the balancing pressure was recorded when water menisci formed on excised vascular bundles (see Supplementary Figure S6 for example). A stereomicroscope (Bausch & Lomb 0.7-3x magnification) was installed above the chamber lid to accurately detect water menisci. Since canopy transpiration differs between night and day conditions and impacts leaf Ψ, leaf Ψ was measured on the same plant when the whole canopy was bagged (i.e., non-transpiring conditions as during night period) or non-bagged (i.e., transpiring conditions as during day period). To do so, we followed the following procedure (see Supplementary Figure S6): The evening prior measurement of leaf Ψ (between 6 PM to 9 PM), two leaves were individually covered in aluminum sleeves (as described above) and subsequently the entire canopy was covered under a black plastic bag. The next day (between 8 AM to 9 AM), the first leaf was excised for measurement of leaf Ψ ‘bagged canopy’. Subsequently, the black plastic bag was removed from the canopy and the second leaf was excised for measurement of leaf Ψ ‘non-bagged canopy’ during the same day (between 12 PM to 2 PM). Iso/anisohydric behavior was determined from slope values of the relationship of leaf Ψ and soil RWC prior to the onset of severe leaf dehydration (Θ_RWC_). Isohydric behavior was defined for slope *a* at *P*>0.05 (i.e., not significantly different from zero), and vice versa for anisohydric behavior.

### Leaf relative water content

Leaf relative water content (leaf RWC) was measured on the same leaf as used for measurement of leaf Ψ ‘bagged canopy’ associated with negligible plant water loss (see Supplementary Figure S6). A 3 cm long bottom portion of the leaf was excised prior measurement of leaf Ψ (as described above), and leaf fresh weight (m_fresh_) was measured using a digital balance (VWR-214B2). Subsequently, the leaf portion was cut into 0.5 cm small pieces and placed into a petri dished filled with water to facilitate full leaf hydration. After 48-h, leaf pieces were gently blotted dry to remove excess surface water and weighed again to obtain leaf saturated weight (m_sat_). Afterwards, leaf pieces were dried in an oven at 70°C temperature for 24-h and measured again to obtain leaf dry weight (m_dry_). Leaf RWC was calculated according to (m_fresh_ – m_dry_) / (m_sat_ – m_dry_) x 100% (Barr and Weatherley, 1962). Subsequently, the onset of severe leaf dehydration (i.e., associated with substantial leaf browning) in response to a decline in soil RWC was determined using a two-segment piecewise linear model and the predicted boundary was defined as Θ_RWC_. The effectiveness of maintaining leaf RWC prior reaching Θ_RWC_ was determined from the steepness of the corresponding slope value.

### Leaf solute potential

Leaf solute potential (leaf Ψ_s_) was measured on the same leaf as used for measurements of leaf Ψ ‘non-bagged canopy’ (see Supplementary Figure S6). A 3 cm long bottom portion of the leaf was excised prior measurement of leaf Ψ, inserted into a 2-ml Eppendorf tube stored on ice (i.e., to minimize solute concentration effects by evaporation), and transferred to a −80°C freezer within 1-h of sampling. For analyses of leaf Ψ_S_, samples (stored for 4-12 weeks) were thawed at room temperature for 15 mins and centrifuged (Eppendorf Centrifuge 5417C) at 11,000 rpm for 14 mins to extract bulk leaf sap. A volume of 10 μL of extracted sap was placed in an osmometer (VAPRO Vapor Pressure Osmometer 5600) and osmolality (in mOsmol kg^-1^) was measured. Subsequently, leaf Ψ_S_ (in MPa) was calculated according [(0.1MPa x ‘osmolality’) / 40.75 mOsmol kg^-1^] (Barrios-Masias et al., 2018). Using linear regression analyses, drought-induced changes in leaf Ψ_S_ prior to Θ_RWC_ were determined from steepness of slope values of the relationship of leaf RWC and soil RWC.

### Leaf stomatal conductance

A porometer (LICOR LI-600) was used to determine stomatal conductance (Gs) of the flag leaf every 1-3 days of each plant (between 12 to 1PM). In the greenhouse, the porometer was equilibrated to the surrounding environment, clamped to the flag leaf, and Gs was recorded when readings stabilized (less than 20 seconds). Severely dehydrated leaves were not evaluated for Gs. A 3-parameter sigmoidal non-linear regression model was fitted to data of Gs and soil RWC to determine the onset of drought-induced reductions in stomatal conductance at 95% of maximum Gs (*Gs_95_*) and soil RWC corresponding to negligible stomatal conductance at 5% of maximum Gs (*Gs_5_*; i.e., full stomatal closure). The difference between *Gs_95_* - *Gs_5_* = *ΔGs_i_* was calculated to quantify the effectiveness of the drought-induced stomatal response. For convenience, we will use the adjectives ‘tight’ and ‘slack’ to refer to a relatively small and large *Gs_i_*, respectively.

### Flag leaf width

Flag leaf width (FLW) was measured every 1-3 days using a digital caliper (Aurora TLV181) at the midpoint of the leaf. Reductions in FLW were assigned to drought-induced leaf rolling (between 1 PM to 2 PM). For each plant, FLW was measured on three leaves and averaged. The extent of leaf rolling prior to the onset of severe leaf dehydration was quantified from the steepness of slope values of the relationship of average FLW and soil RWC (i.e., slope of zero at *P*>0.05 = no leaf rolling).

### Kernel weight

Wheat spikes from plants which did not have flag leaves harvested were harvested 20 days to allow for the control plant seeds to mature after the end of experiment 1 and 2. Excised wheat spikes were placed in paper bags and transported to the lab within 2-h. Spikes were threshed using a hand thresher and counted. Following the procedure of Qu et al. (2022), the TKW of the plants was measured by counting 100 seeds, weighing them (Ohaus TP600S) and then multiplying by 10. In the cases where the plants did not produce 250 seeds, the weight per seed was determined and multiplied by 1,000 to get the TKW.

### Data analyses

Data was analyzed using RStudio (version 2022.02.3+492). Sigmaplot (Version 8.0) was used to create graphs and for the regressions (PLR and linear regressions). Slope values for the PLR model (leaf RWC) was identified before (slope *a*) and after (slope *b*) Θ_RWC_. A linear regression was run on the leaf Ψ_S_, FLW, transpiring and non-transpiring leaf Ψ data prior to Θ_RWC_ and slope c was determined. A linear mixed effects model was used to analyze leaf RWC, leaf Ψ_S_, FLW, transpiring and non-transpiring leaf Ψ. A ‘cultivar’ fixed factor and ‘individual’ random factor (to account for repeated measures) was used with soil RWC as the covariate in the model. F and p-values were found for each parameter using a Type III ANOVA. Statistically significant differences in means were defined as *P*<0.05. If interaction effects were not found to be significant, a Type II ANOVA was conducted. If any significance was found, a post-hoc Tukey’s t-test was run to determine which differences in means were statistically significant. A Type III ANOVA was also conducted for the TKW data to determine if ‘cultivar’, ‘treatment’ (soil RWC group), or an interaction effect existed. A post-hoc Tukey’s analysis was done after to determine which groups differed statistically. Statistical significance was defined as *P*<0.05.

## RESULTS

Compared to ‘Stettler’ and ‘AAC Viewfield’, ‘Superb’ successfully maintained leaf RWC and to lower levels of soil RWC under progress drought stress (Figure 1). For ‘Superb’ (Figure 1A), Θ_RWC_ corresponded to a soil RWC of 36% and no significant change in leaf RWC was detected during leaf hydration phase I (slope *a* at *P*=0.76). In contrast, for ‘Stettler’ (Figure 1B) and ‘AAC Viewfield’ (Figure 1C), Θ_RWC_ was reached at a higher soil RWC of 48% and 51%, respectively, and slope a was *P*≤ 0.05 during leaf hydration phase I. During phase I, there was a significant ‘cultivar’ effect (*P*=*0.04*, Supplementary Table S3) with a statistically significant difference between ‘Superb’ and ‘AAC Viewfield’ (P=0.02, Supplementary Table S4). Following Θ_RWC_, severe leaf dehydration and a substantial drop in leaf RWC was detected in all three CWRS wheat cultivars. For ‘Superb’ (Figure 1A), this reduction was statistically significant (slope *b* at *P*<0.01). For ‘Stettler’ and ‘AAC Viewfield’, slope *b* (*P*>0.05) was shallower (Figure 1B and 1C).

**Figure 1.**
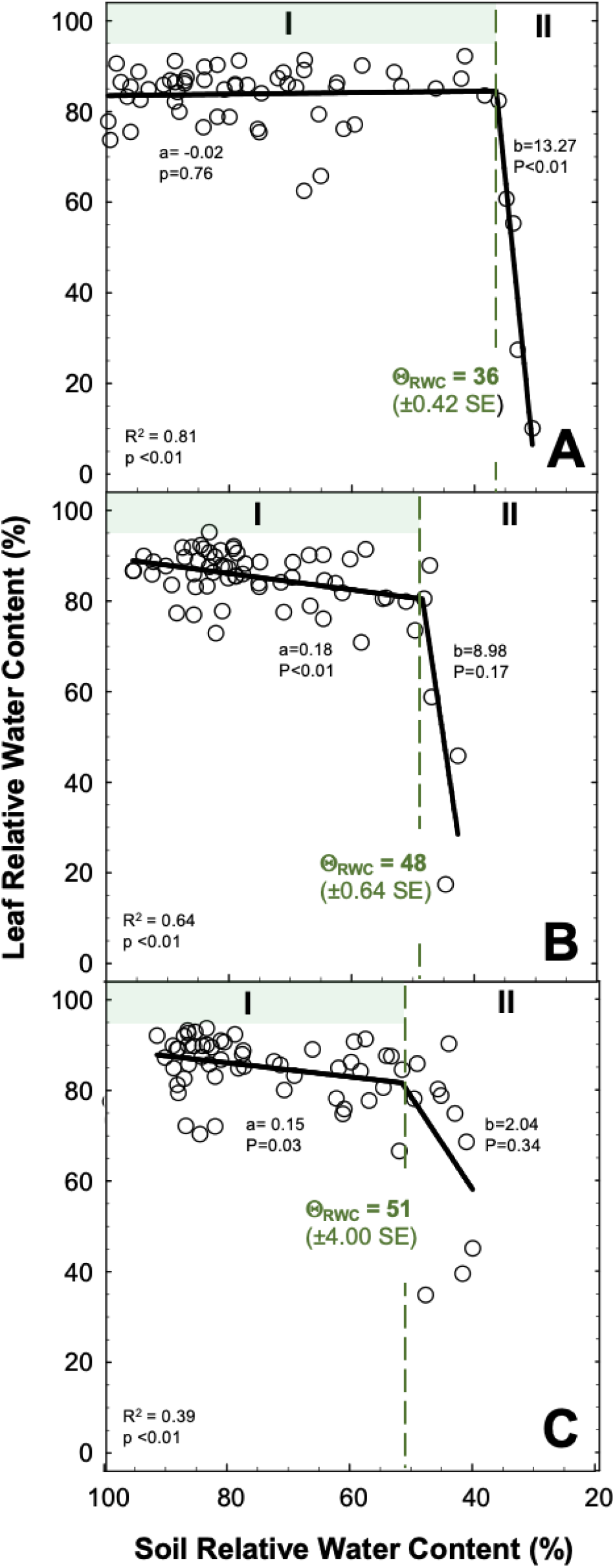
Effect of water stress by drought on leaf relative water content of three wheat cultivars (panel A ‘Superb’, panel B ‘Stettler’, panel C ‘AAC Viewfield’). Each symbol represents a single measurement of an individual plant. The solid line is a two-segment piecewise linear regression model fitted across data points (for R^2^ and p values, see bottom left in each panel); the vertical dashed line indicates the predicted threshold of onset of severe leaf dehydration (Θ_RWC_). Roman numerals I and II indicate the corresponding phases of leaf hydration status prior and after Θ_RWC_. Parameters a and b are slope values of phase I and II, respectively, with corresponding p-values and coefficient of determination (R^2^).

During leaf hydration phase I, leaf Ψ was differentially affect under non-transpiring and transpiring conditions when soil RWC declined (Figure 2). For a bagged (non-transpiring) canopy (Figures 2A to 2C) leaf Ψ was maintained at approximately −0.85 MPa, −1 MPa, and 0.7 MPa for ‘Superb’, ‘Stettler’, and ‘AAC Viewfield’, respectively. Reductions in leaf Ψ in response to a decline in soil RWC were negligible in all three CWRS wheat cultivars and corresponding slope *c* was at *P*>0.05. Analysis of the differences in means indicated a significant ‘cultivar’ effect was found for non-transpiring canopy in phase I (Supplementary Table S5). The means of ‘AAC Viewfield’ and ‘Stettler’ were found to be statistically different (Supplementary Table S6). For a transpiring canopy (Figures 2D to 2F), a significant decline in leaf Ψ with soil RWC was detected in all three CWRS wheat cultivars (slope *c* at *P*<0.05). For a transpiring canopy, there was a significant ‘cultivar’ effect in phase I (Supplementary Table S5) and a statistically significant difference in the mean was predicted between ‘AAC Viewfield’ and ‘Superb’ (Supplementary Table S7).

**Figure 2.**
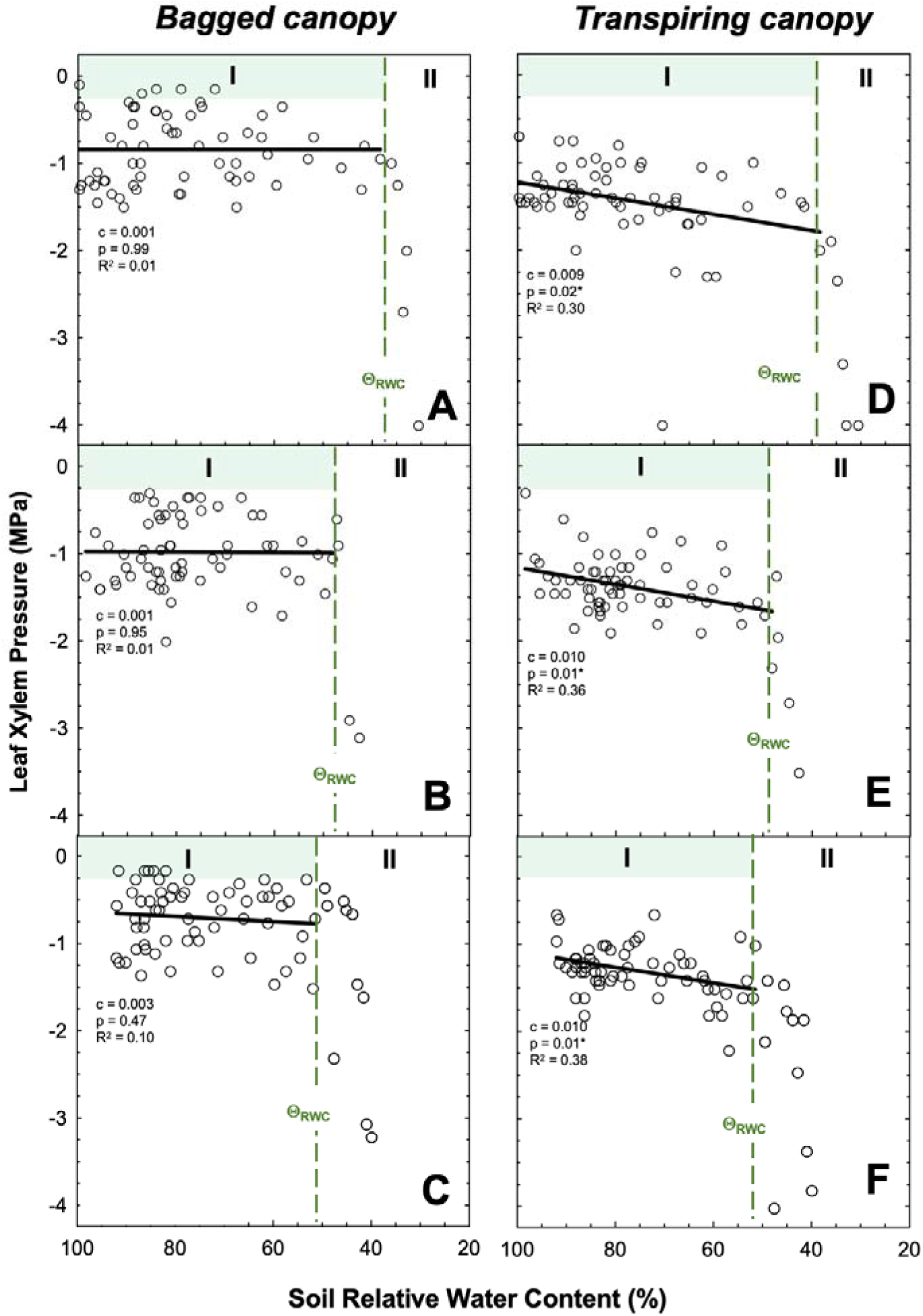
Effect of water stress by drought on leaf xylem pressure (leaf Ψ) under non-transpiring (panels A to C) and transpiring (panels D to F) of three wheat cultivars (panels A/D ‘Superb’, panels B/E ‘Stettler’, panels C/F ‘AAC Viewfield’). Each symbol represents a single measurement of an individual plant. The solid line is a simple linear regression fitted across datapoints of phase I; parameter *c* is the slope value with corresponding p-values and coefficient of determination (R^2^). The vertical dashed line is the superimposed threshold Θ_RWC_ (see Figure 1). Roman numerals I and II indicate the corresponding phases of leaf hydration status prior and after Θ_RWC_.

Leaf Ψ_S_ remained at a similar level (approximately 1.15 MPa) in response to reductions in soil RWC during phase I for ‘Superb’ (slope c = 0.001, *P*>0.05, Figure 3A) and ‘Stettler’ (slope *c* = −0.001, *P*>0.05 Figure 3B). For ‘AAC Viewfield’, our data indicate a slight reduction in leaf Ψ_S_ (from −1.05 to −1.25 MPa) during phase I (slope *c* = 0.004, *P*<0.05, Figure 3C). Further analysis showed that there was no predicted no significant ‘cultivar’ effect in phase I (Supplementary Table S5). Following Θ_RWC_, leaf Ψ_S_ dropped substantially to more negative values in all three CWRS wheat cultivars (Figures 3A to 3C).

**Figure 3.**
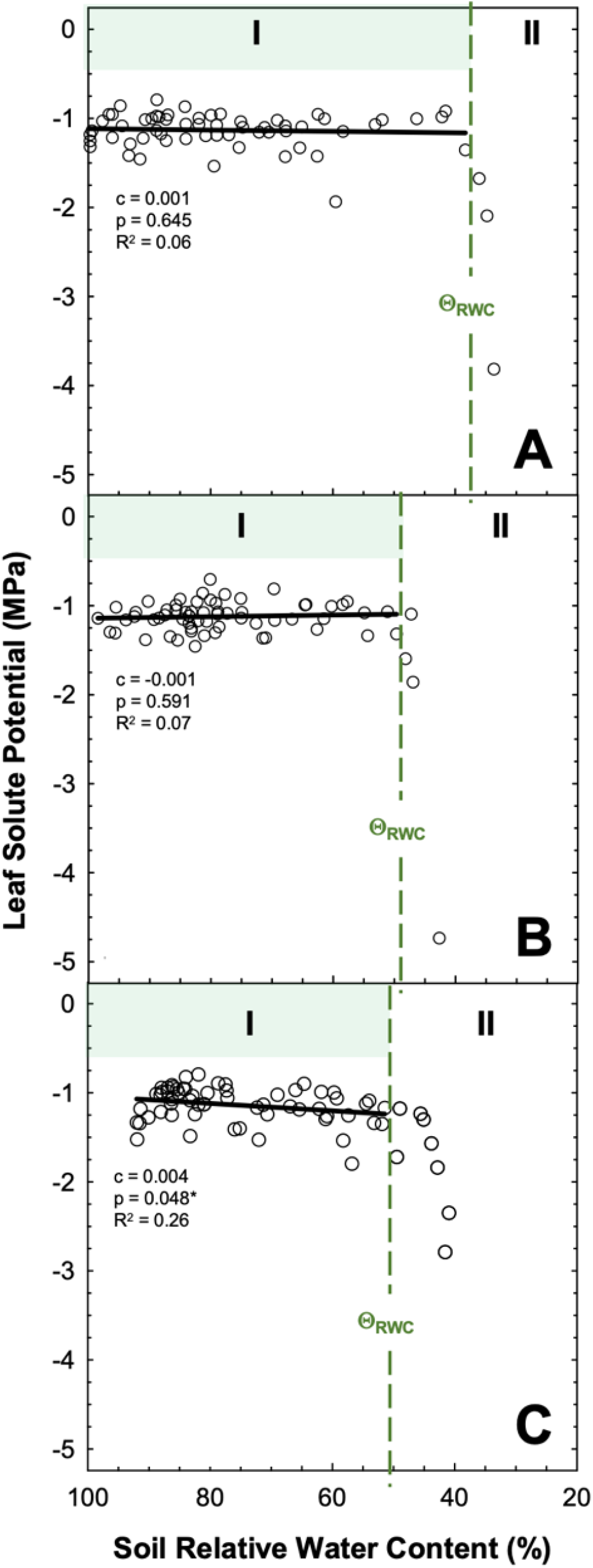
Effect of water stress by drought on leaf solute potential of three wheat cultivars (panel A ‘Superb’, panel B ‘Stettler’, panel C ‘AAC Viewfield’). Each symbol represents a single measurement of an individual plant. The solid line is a simple linear regression fitted across datapoints of phase I; parameter *c* is the slope value with corresponding p-values and coefficient of determination (R^2^). The vertical dashed line is the superimposed threshold Θ_RWC_ (see Figure 1). Roman numerals I and II indicate the corresponding phases of leaf hydration status prior and after Θ_RWC_.

Stomatal regulation in response to soil RWC differed among the three cultivars (Figure 4). For ‘Superb’ (Figure 4A), Gs_95_ and Gs_5_ corresponded to a soil RWC of 72% and 48% respectively, resulting in a ΔGs_i_ of 24%. For ‘Stettler’ (Figure 4B), Gs_95_ and Gs_5_ were 82% and 46%, respectively, giving a ΔGs_i_ of 36%. For ‘AAC Viewfield’ (Figure 4C), Gs_95_ and Gs_5_ were at 99% and 38%, respectively, and ΔGs_i_ was 61%. In addition, Gs_5_ was reached prior to Θ_RWC_ in ‘Superb’ (Figure 4A). On the other hand, Gs5 coincided with Θ_RWC_ in ‘Stettler’ and occurred after Θ_RWC_ in ‘AAC Viewfield’ (Figures 4B and 4C).

**Figure 4.**
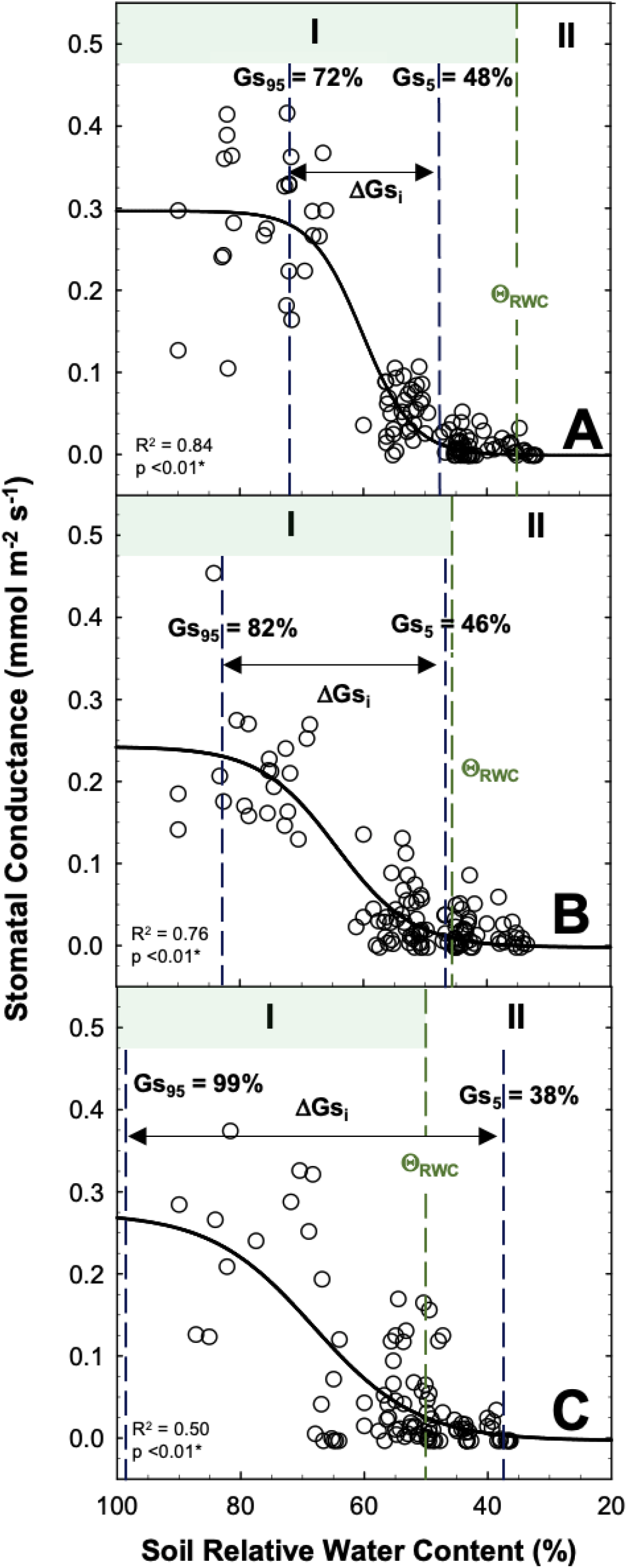
Relation of soil relative water content and stomatal conductance of three wheat cultivars (panel A ‘Superb’, panel B ‘Stettler’, panel C ‘AAC Viewfield’). Each symbol represents a single measurement on an individual plant. The solid line is a non-linear sigmoidal three-parameter regression fitted across datapoints. The vertical blue dashed linear are the thresholds of 95% (Gs_95_, onset of stomatal closure) and 5% (Gs_5_, full stomatal closure) of maximal stomatal. Parameter ΔGs_i_ is the difference between Gs_95_ and Gs_5_. The green colored dashed line is the superimposed threshold Θ_RWC_ (see Figure 1). Roman numerals I and II indicate the corresponding phases of leaf hydration status prior and after Θ_RWC_.

During phase I, a decline in FLW by leaf rolling was detected in ‘Superb’ but not for ‘Stettler’ and ‘AAC Viewfield’ (Figure 5). For ‘Superb’ (Figure 5A), FLW declined significantly from on average 16 to 10 mm at Θ_RWC_ (slope *c* = 0.138, *P*<0.05). For ‘Stettler’ (c = 0.0024, *P*>0.05 Figure 5B) and ‘AAC Viewfield’ (c = 0.0013, *P*>0.05, Figure 5C), regression analyses predict no significant reduction in FLW during phase I. The FLW for both ‘Stettler’ and ‘AAC Viewfield’ was maintained at approximately 13 mm in phase I. Statistical analyses predicted a ‘cultivar’ effect during phase I (Supplementary Table S3), and a statistically significant difference in means was determined between ‘AAC Viewfield’ and ‘Superb’ (*P*<0.05; Supplementary Table S8).

**Figure 5.**
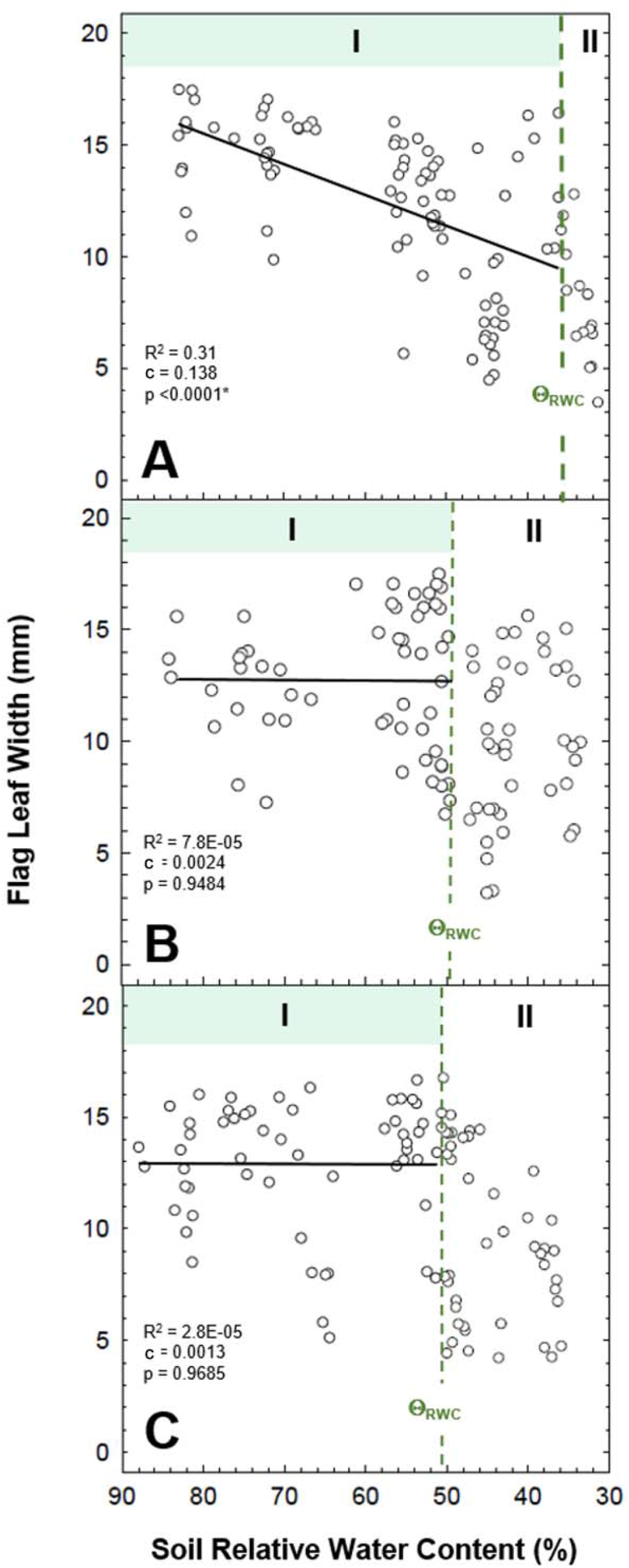
Effect of water stress by drought on flag leaf width of three wheat cultivars (panel A ‘Superb’, panel B ‘Stettler’, panel C ‘AAC Viewfield’). Each symbol represents a single measurement of an individual plant. The solid line is a simple linear regression fitted across datapoints of phase I; parameter c is the slope value with corresponding p-values and coefficient of determination (R^2^). The vertical dashed line is the superimposed threshold Θ_RWC_ (see Figure 1). Roman numerals I and II indicate the corresponding phases of leaf hydration status prior and after Θ_RWC_.

TKW was maintained during phase I in all three cultivars, and reductions in TKW occurred when threshold Θ_RWC_ was passed (Figure 6). For ‘Superb’, a significant reduction in TKW from 33 to 9 g was detected at soil RWC of <45% that is during phase II following Θ_RWC_ (Figure 6A). In ‘Stettler’, TKW dropped significantly at <45% soil RWC that followed Θ_RWC_ of 48% from 35 to 6 g (Figure 6B). In ‘AAC Viewfield’, TKW dropped significantly at <60% soil RWC and, again, after reaching Θ_RWC_ of 58% from 36 to 13 g (Figure 6C). In general, ‘Superb’ exhibited better TKW compared to ‘Stettler’ and ‘AAC Viewfield’ under normal soil moisture (90%). Statistical analyses supported differences between cultivars and treatments (Supplementary Table S9). Differences in means for treatments were statistically significant (Supplementary Table S10). Moreover, the means of ‘AAC Viewfield’ and ‘Superb’ were statistically different (Supplementary Table S11).

**Figure 6.**
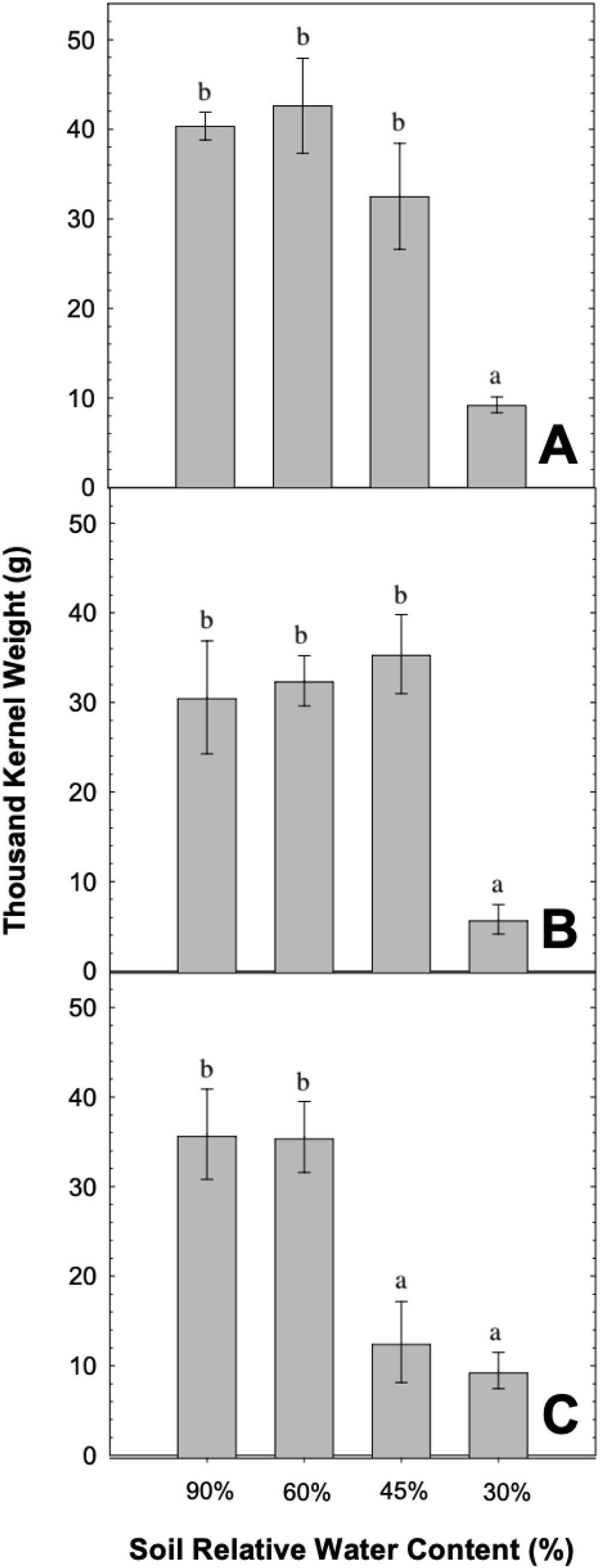
Thousand kernel weight at different levels of soil RWC of three wheat cultivars (panel A ‘Superb’, panel B ‘Stettler’, panel C ‘AAC Viewfield’). Each bar is the mean ± standard error of mean of n=3 plants. Different letter indicates statistically significant differences at *P*<0.05 (posthoc Tukey test).

Soil core samples were extracted to provide an estimation of soil water heterogeneity and consequently root water uptake. A statistical analysis of the means of the cores shows that there are no statistically significant differences between cultivar and soil depths (Appendices 6 and 7). The soil GWC in ‘Superb’ starts between 8% to 16% for well-watered plants above 90% soil RWC at all three depths. By the time ‘Superb’ hit 30% soil RWC, the soil GWC was close to 0% at all three depths (Appendix 5B). Similarly, ‘Stettler’ starts between 6% and 19% soil GWC for well-watered plants and hits close to 0% soil GWC at approximately 35% soil RWC (Appendix 5C). ‘AAC Viewfield’ followed a similar trend where it started between 4% and 17% soil GWC for well-watered plants while it was close to 0% soil GWC for plants at approximately 35% soil RWC (Appendix 5D). While there is a general trend downwards in soil GWC of soil cores from top to bottom of pot these changes are not statistically significant (Appendices 6 and 7). Furthermore, representative pictures of root system did not indicate substantial differences in maximum rooting depth between the three CWRS wheat cultivars (Supplementary Figure S7). The roots of each cultivar surpassed the depth of their pots (60 cm).

## DISCUSSION

In this study we show that CWRS wheat cultivars ‘Superb’, ‘Stettler’, and ‘AAC Viewfield’ exhibit negligible OA in response to drought and that sustained kernel development in these cultivars is linked to ‘tight’ stomatal regulation and effective leaf rolling. Although all three CWRS wheat cultivars exhibited anisohydric behavior under transpiring canopy conditions, we found that this behavior was not sufficient to trigger OA. The onset of reductions in TKW was closely linked to threshold Θ_RWC_, which suggests Θ_RWC_ as a predictor of yield losses under drought conditions. Compared to ‘Stettler’ and ‘AAC Viewfield’, ‘Superb’ exhibiting tight stomatal regulation and effective leaf rolling during phase I was most effective in maintaining TKW and as reflected by its Θ_RWC_ thresholds at a relative low soil RWC of 36%. While the slopes in phase II all decreased after Θ_RWC_ there was no statistical support for the slopes for ‘Stettler’ and ‘AAC Viewfield’ which can most likely be explained by the lack of data points after the threshold. Regardless, our study highlights physiological characteristics that may improve drought tolerance when used as future breeding targets for CWRS wheat cultivars. An overview of measured cultivar-specific physiological traits is given in Table 2 that may serve as a reference for future studies into drought tolerance of CWRS wheat cultivars.

Leaf OA minimizes the extend of drought-induced cell dehydration due to its positive effect on lower cell water potential and is typically the result of organic solutes accumulating inside the cell that do not interfere with enzyme activities even at high concentrations (Turner and Jones, 1980; Turner, 1986; Kramer and Boyer, 1995). No significant reduction in leaf Ψ_S_ was observed in ‘Superb’ and ‘Stettler’ until reaching Θ_RWC_. This suggests that no OA occurs in both CWRS wheat cultivars as the leaf Ψ_S_ does not decrease and therefore no accumulation of solutes occurs while leaf RWC is relatively stable during phase I. We speculate that the slight, but significant, reduction in leaf Ψ_S_ of ~0.05 MPa for ‘AAC Viewfield’ was most likely linked to the reduction in leaf RWC until reaching Θ_RWC_ and a solute concentration effect (Figure 1). Interestingly, for ‘Superb’ leaf RWC was maintained during phase I even in the absence of OA which points to other physiological mechanisms supporting this behavior. Moreover, the method described in Babu et al. (1999) was used to evaluate OA. The solute accumulation effect under decreasing soil RWC was estimated by measured leaf Ψ_S_ minus leaf Ψ_S0_ (leaf Ψ_S_ due to concentration effect) at 80% soil RWC. In all three CWRS wheat cultivars, this method predicted a relatively small solute accumulation effect – if any (‘Superb’ at −0.10 MPa; ‘Stettler’ at −0.18 MPa and ‘AAC Viewfield’ at −0.20 MPa) supporting the previous findings (Supplementary Figure S8). Overall, active solute accumulation (i.e., OA) played a negligible role in all three CWRS wheat cultivars.

Drought induced stomatal closure is a mechanism that prevents excessive plant water loss (Tyree and Sperry, 1988; Brodribb and McAdam, 2011). Our data point to cultivar-specific differences in stomatal regulation with respect to Gs_95_ and Gs_5_. Stomata started closing (Gs_95_) the earliest in ‘AAC Viewfield’. Meanwhile, both ‘Superb’ and ‘Stettler’ had a much later initiation of stomatal closure, so the plants were able to transpire fully for longer than ‘AAC Viewfield’. Although this was the case, the stomata did close much earlier (Gs_5_) in ‘Superb’ and ‘Stettler’ than it did in ‘AAC Viewfield’. For ‘Superb’, the margin of Gs_95_ (onset of stomatal closure) to Gs_5_ (stomatal closure) was relatively small (i.e., soil RWC from 72% to 48%), which indicates a relatively ‘tight’ stomata control. In contrast, our data for ‘AAC Viewfield’ exhibited the opposite response where stomatal closure was initiated early on (Gs_95_) when soil RWC declined and Gs_5_ was reached at soil RWC of 38%. This relatively ‘slack’ stomatal control may be interpreted as a sign of drought intolerance from a physiological perspective because the plant does not adjust its stomata quickly in response to drying and continues to transpire (Tardieu et al., 2018). ‘Stettler’ played an intermediate role but behaved more similarly to ‘Superb’. Our data suggest that a ‘tight’ stomata control under drought and prior reaching Θ_RWC_ is linked to maintenance of leaf RWC under drought in the absence of OA. On the other hand, relative late onset of stomata closure in Superb can be interpret as a beneficial trait to maintain leaf gas exchange (i.e., CO_2_ assimilation) under water limiting conditions that has positive impacts on grain development. However, our data highlight that cultivar-specific stomatal regulation in response to progressive drought did not affect the degree of isohydricity. For example, although ‘Superb’ and ‘AAC Viewfield’ differed substantial in their stomatal response they showed both showed anisohydric behavior under transpiring canopy conditions. This suggest that anisohydric behavior associated with reduction in leaf Ψ does not trigger OA in CWRS wheat cultivars.

Leaf rolling has several benefits for maintaining hydration status including minimizing transpirational water loss while stomata can remain in an open state and protecting the leaf from high radiation (Kadiogiu and Terzi, 2007). Our data on drought-induced changes in FLW by leaf rolling indicate that leaf rolling happens prior to Θ_RWC_ in ‘Superb’ but not in ‘Stettler’ and ‘AAC Viewfield’. This suggests that leaf rolling in ‘Superb’ plays a part in maintaining its leaf RWC. Moreover, ‘Superb’ has the lowest Gs_95_ of the three cultivars, which could be due to the leaf rolling seen in ‘Superb’ but not in the other two cultivars. Visual approaches or sustained yield under drought have been used by breeders to select breeding material with improved drought tolerance. This is so because they can be much faster and cheaper compared to physiological tests (Jones, 1978). In this context our data indicate that leaf rolling, or observation of leaf rolling are not necessarily a sign of drought stress in CWRS wheat but can acts as a mechanism providing for improved drought tolerance in specific cultivars.

The homogeneity of soil water in deep pots such as the one used in this project plays a big part when conducting drought studies. Rewatering plants from the top causes water to remain in the top of the pot (Turner, 2019) which could make estimates of soil RWC inaccurate as they are taken as an average of the pot. Our data indicate that there was no statistical difference between soil GWC at different depths suggesting that possible gradient in soil moisture along the pot profile were negligible. Furthermore, all three cultivars grew root systems that reached the bottom soil layer of pots indicating that roots had access to all soil layers (Supplementary Figure S5). This falls in line with previous research as wheat roots stop growing after the heading stage (Hurd, 1964) so the wheat roots are already established and at the bottom of the pot by the time the drydown was applied. This suggests that the differences in drought performance seen in the cultivars were not due to rooting depth. Nevertheless, this highlights that further research on ‘Superb’, ‘Stettler’, and ‘AAC Viewfield’ should target a detailed investigation of root physiological responses and root architecture to drought.

Wheat TKW weight decreases as soils become drier in part due to the lack of water availability causing plants to form grains in a shortened time frame (Poudel et al., 2020). Similar results were found in this study on CWRS wheat cultivars. However, the reduction in TKW seen between cultivars was closely linked to Θ_RWC_. In turn, this points towards the importance of this threshold as a predictor for TKW for the cultivars under water limited conditions. Leaf RWC provides information on plant water status (Mullan and Pertragalla, 2012). In turn, a reduction in leaf RWC is linked to reductions in yield and TKW (Schofeld et al., 1988; Tahara et al., 1990). Consequently, an CWRS wheat cultivar with a lower Θ_RWC_ would perform better in drought as it would be able to manage itself to conserve water. CWRS wheat cultivars with tighter stomatal control (‘Superb’ and ‘Stettler’) close their stomata quickly when water availability decreases (Busk et al., 1999) thereby decreasing transpiration and preventing severe leaf dehydration (Tardieu et al., 2018).

**Table 1.** General overview of leaf physiological traits observed in this study for three wheat cultivars (‘Superb’, ‘Stettler’, and ‘AAC Viewfield’) under water stress by drought during phase I (prior to reaching Θ_RWC_) Signs indicate relative differences in characteristics between cultivars; ‘better’ (+), ‘same’ (o), or ‘absent’ (-)

| Response Trait | Cultivar |  |  |
| --- | --- | --- | --- |
|  | *Superb* | *Stettler* | *AAC Viewfield* |
| Relative water content maintenance | + | - | - |
| Iso-/aniso-hydric behavior | o | o | o |
| Osmotic adjustment | - | - | - |
| Stomatal behavior | + | o | - |
| Leaf rolling | + | - | - |

## ACKNOWLEDGEMENTS

We would like to thank Dr. Pierre Hucl (University of Saskatchewan) for assisting with cultivar selection, Dr. Megan Bartlett (University of California, Davis) and Dr. Penny Tricker (University of Adelaide) for important feedback on plant physiological behaviors. This work was supported by UBC-LFS start funds provided to T.K. and G.S.B.

## REFERENCES

Bagatta, M., Pacifico, D., & Mandolino, G. (2008). Evaluation of the osmotic adjustment response within the genus Beta. Journal of Sugar Beet Research, 45(3/4), 119.

Barr, H. D., & Weatherley, P. E. (1962). A re-examination of the relative turgidity technique for estimating water deficit in leaves. Australian Journal of Biological Science, 15(3), 413–428.

Blum, A. (2017). Osmotic adjustment is a prime drought stress adaptive engine in support of plant production. Plant, cell & environment, 40(1), 4–10.

Blum, A., & Pnuel, Y. (1990). Physiological attributes associated with drought resistance of wheat cultivars in a Mediterranean environment. Australian Journal of Agricultural Research, 41(5), 799–810.

Blum, A., Zhang, J., & Nguyen, H. T. (1999). Consistent differences among wheat cultivars in osmotic adjustment and their relationship to plant production. Field Crops Research, 64(3), 287–291.

Brodribb, T. J., & McAdam, S. A. (2011). Passive origins of stomatal control in vascular plants. Science, 331(6017), 582–585.

Busk, P. K., Borrell, A., Kizis, D., & Pagès, M. (1999). Abscisic acid perception and transduction. In Biochemistry and Molecular Biology of Plant Hormones (pp. 491–512). Elsevier BV.

Cai, T., Meng, X., Liu, X., Liu, T., Wang, H., Jia, Z., … & Ren, X. (2018). Exogenous hormonal application regulates the occurrence of wheat tillers by changing endogenous hormones. Frontiers in plant science, 9, 1886.

Campbell, C. A., Zentner, R. P., Basnyat, P., Wang, H., Selles, F., McConkey, B. G., … & Cutforth, H. W. (2007). Water use efficiency and water and nitrate distribution in soil in the semiarid prairie: Effect of crop type over 21 years. Canadian Journal of Plant Science, 87(4), 815–827.

Curtis, T., & Halford, N. G. (2014). Food security: the challenge of increasing wheat yield and the importance of not compromising food safety. The Annals of applied biology, 164(3), 354–372.

Cuthbert, R. D., DePauw, R. M., Knox, R. E., Singh, A. K., McCallum, B., & Fetch, T. (2018). AAC Viewfield hard red spring wheat. Canadian Journal of Plant Science, 99(1), 102–110.

DePauw, R. M., Knox, R. E., Clarke, F. R., Clarke, J. M., & McCaig, T. N. (2009). Stettler hard red spring wheat. Canadian Journal of Plant Science, 89(5), 945–951.

De Jong, R., Campbell, C. A., Zentner, R. P., Basnyat, P., Cutforth, H., & Desjardins, R. (2008). Quantifying soil water conservation in the semiarid region of Saskatchewan, Canada: Effect of fallow frequency and N fertilizer. Canadian Journal of Soil Science, 88(4), 461–475.

Henson, I. E., Jensen, C. R., & Turner, N. C. (1989). Leaf gas exchange and water relations of lupins and wheat. I. Shoot responses to soil water deficits. Functional Plant Biology, 16(5), 401–413.

Hurd, E. A. (1964). Root study of three wheat varieties and their resistance to drought and damage by soil cracking. Canadian journal of plant science, 44(3), 240–248.

Innes, P., & Blackwell, R. D. (1981). The effect of drought on the water use and yield of two spring wheat genotypes. The Journal of Agricultural Science, 96(3), 603–610.

Jones, H. G. (1978). Modelling diurnal trends of leaf water potential in transpiring wheat. Journal of Applied Ecology, 10(2), 613–626.

Jones, B. C. (2008). Microclimate conditions within the rolled leaves of Lepanthes helleri (Orchidaceae). Tropical Ecology and Conservation [Monteverde Institute]. 260.

Kadioglu, A., & Terzi, R. (2007). A dehydration avoidance mechanism: Leaf rolling. [Das Einrollen von Blättern als Schutz vor Austrocknung] The Botanical Review, 73(4), 290–302.

Knipfer, T., Reyes, C., Momayyezi, M., Brown, P. J., Kluepfel, D., & McElrone, A. J. (2020) A comparative study on physiological responses to drought in walnut genotypes (RX1, Vlach, VX211) commercially available as rootstocks. Trees, 34, 665–678.

Kramer, P. J., & Boyer, J. S. (1995). Water relations of plants and soils. Academic press.

Mahmood, T., Abdullah, M., Ahmar, S., Yasir, M., Iqbal, M. S., Yasir, M., Ur Rehman, S., Ahmed, S., Rana, R. M., Ghafoor, A., Nawaz Shah, M. K., Du, X., & Mora-Poblete, F. (2020). Incredible Role of Osmotic Adjustment in Grain Yield Sustainability under Water Scarcity Conditions in Wheat (Triticum aestivum L.). Plants (Basel, Switzerland), 9(9), 1208.

Morgan, J. M. (1977). Differences in osmoregulation between wheat genotypes. Nature, 270(5634), 234–235.

Morgan, J. M. (1980). Osmotic adjustment in the spikelets and leaves of wheat. Journal of Experimental Botany, 31(2), 655–665.

Morgan, J. M., & Condon, A. G. (1986). Water use, grain yield, and osmoregulation in wheat. Functional plant biology, 13(4), 523–532.

Mullan, D., & Pietragalla, J. (2012). Leaf relative water content. Physiological breeding II: A field guide to wheat phenotyping, 25–27.

Oppenheimer, H. R. (1960). Adaptation to drought: xerophytism. Plant-water relationships in arid and semi-arid conditions: reviews of research.

Poudel, M. R., Ghimire, S., Dhakal, K. H., Thapa, D. B., & Poudel, H. K. (2020). Evaluation of wheat genotypes under irrigated, heat stress and drought conditions. Journal of Biology and Today’s World, 9(1), 1–12.

Qu, X., Li, C., Liu, H., Liu, J., Luo, W., Xu, Q., … & Ma, J. (2022). Quick mapping and characterization of a co-located kernel length and thousand-kernel weight-related QTL in wheat. Theoretical and Applied Genetics, 135(8), 2849–2860.

Tahara, M., Carver, B. F., Johnson, R. C., & Smith, E. L. (1990). Relationship between relative water content during reproductive development and winter wheat grain yield. Euphytica, 49(3), 255–262.

Tardieu, F., Simonneau, T., & Muller, B. (2018). The physiological basis of drought tolerance in crop plants: A scenario-dependent probabilistic approach. Annual Review of Plant Biology, 69(1), 733–759.

Townley-Smith, T. F., Humphreys, D. G., Czarnecki, E., Lukow, O. M., McCallum, B. M., Fetch, T. G., … & Brown, P. D. (2010). Superb hard red spring wheat. Canadian Journal of Plant Science, 90(3), 347–352.

Turner, N. C., & Jones, H. G. (1980). Turgor maintenance by osmotic adjustment. A review and evaluation. Adaptation of plants to water and high temperature stress., 87–103.

Turner, N. C. (1981). Techniques and experimental approaches for the measurement of plant water status. Plant and Soil, 58(1/3), 339–366.

Turner, N. C. (1986). Crop water deficits: a decade of progress. Advances in agronomy, 39, 1–51.

Turner, N. C. (2017). Turgor maintenance by osmotic adjustment, an adaptive mechanism for coping with plant water deficits: Turgor maintenance by osmotic adjustment. Plant, Cell and Environment, 40(1), 1–3.

Turner, N. C. (2019). Imposing and maintaining soil water deficits in drought studies in pots. Plant and Soil, 439, 45–55.

Trillo, N., Fernandez, R.J. (2005). Wheat Plant Hydraulic Properties Under Prolonged Experimental Drought: Stronger Decline in Root-system Conductance than in Leaf Area. Plant Soil, 277, 277–284.

Tyree, M. T., & Sperry, J. S. (1988). Do woody plants operate near the point of catastrophic xylem dysfunction caused by dynamic water stress? Answers from a model. Plant physiology, 88(3), 574–580.

Willick, I. R., Lahlali, R., Vijayan, P., Muir, D., Karunakaran, C., & Tanino, K. K. (2018). Wheat flag leaf epicuticular wax morphology and composition in response to moderate drought stress are revealed by SEM, FTIR-ATR and synchrotron X-ray spectroscopy. Physiologia plantarum, 162(3), 316–332.

Xoconostle-Cazares, B., Ramirez-Ortega, F. A., Flores-Elenes, L., & Ruiz-Medrano, R. (2010). Drought tolerance in crop plants. American journal of plant physiology, 5(5), 241–256.

