## Supplementary Figures and Tables for "Leaf hydration status under drought is predominantly linked to stomatal regulation and leaf roll but not osmotic adjustment in Canadian hard red spring wheat (*Triticum aestivum*) cultivars"

### Supplementary Material

**Supplementary Figure S1.** Daily temperature (panels A and D), relative humidity (panels B and E), and VPD (panels C and F) during experiments 1 (panels A-C, *June to September 2021*) and 2 (panels D-F, *February to June 2022*). The drydown is indicated by the shaded blue region. The imposed red line represents when the plants were harvested.

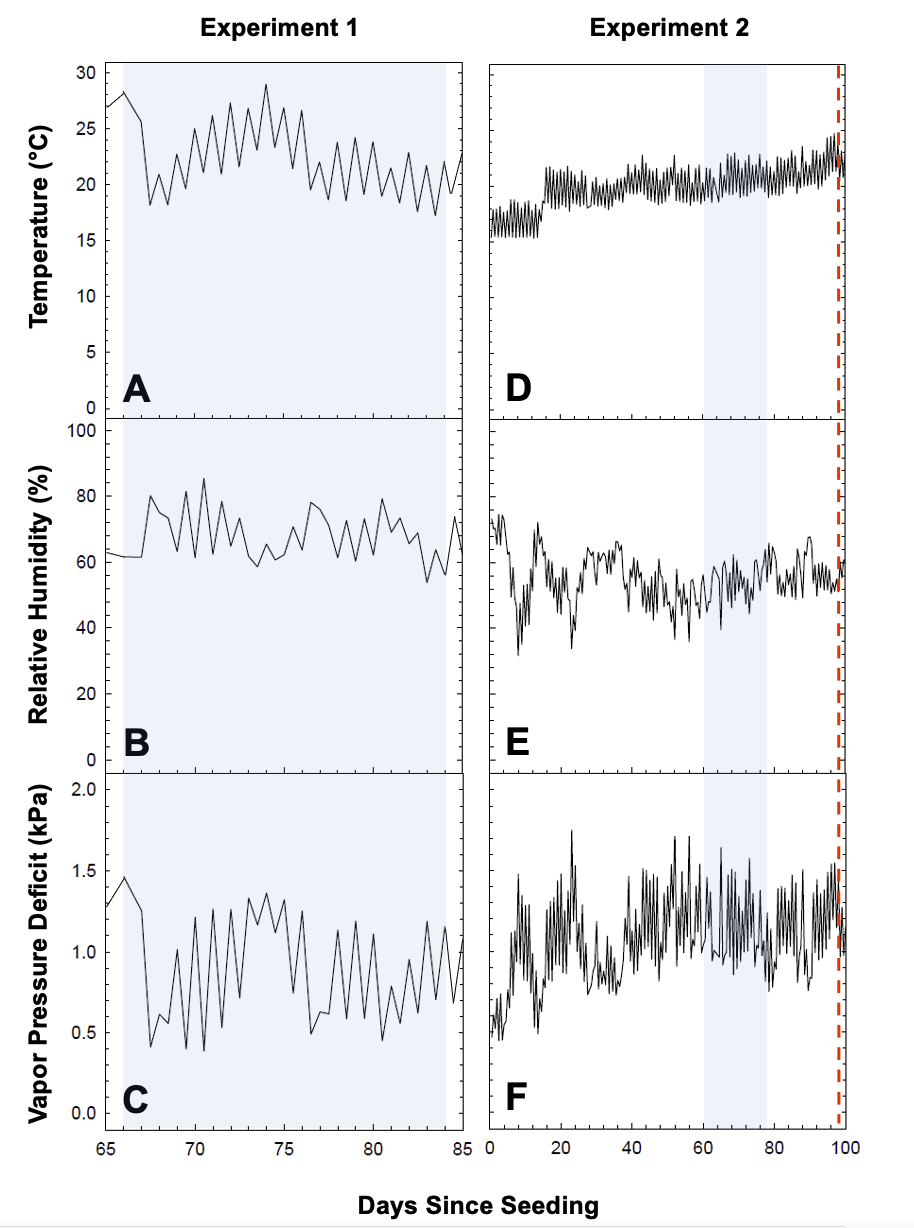

**Supplementary Figure S2.** Experimental randomized block design for (panel A) experiment 1 and (panel B) experiment 2 including cultivars and treatments.

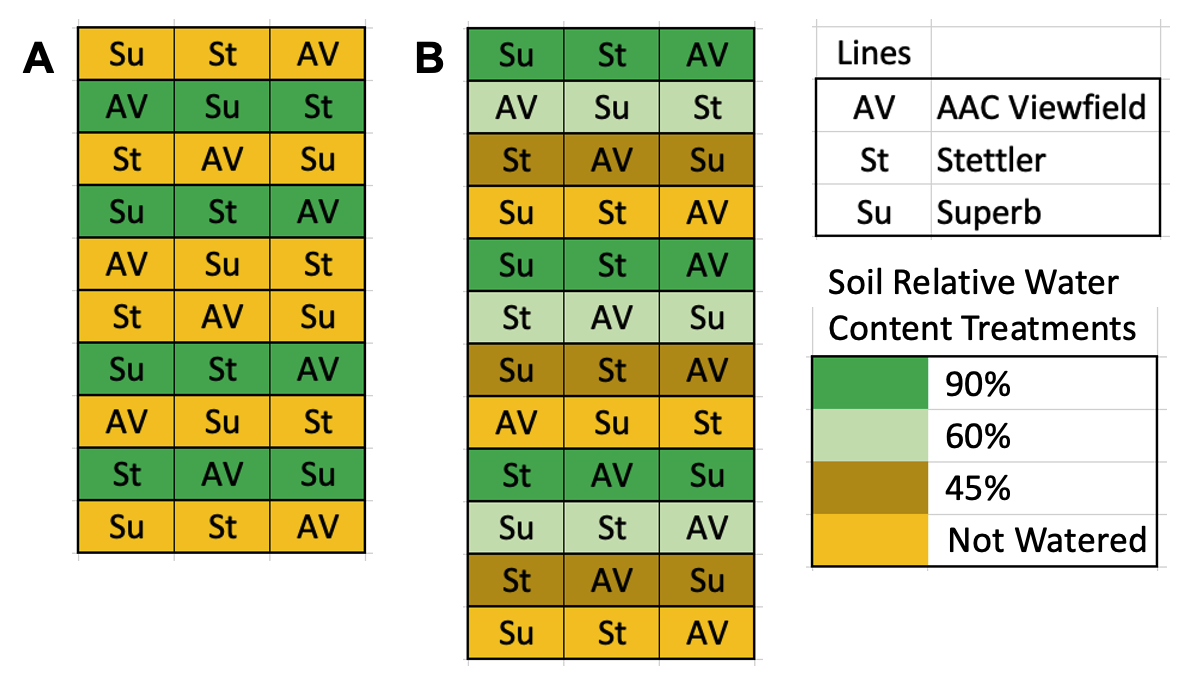

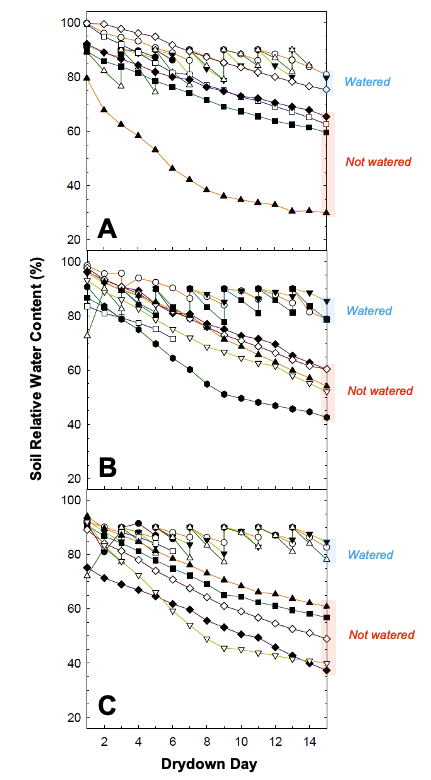
**Supplementary Figure S3.** Temporal changes of soil relative water content of plants measured for experiment 1 (Figure 1 to 5). Watered plants (90% soil relative water content) and not watered plants (water withheld) are shown by the blue and red shaded regions, respectively. Each plant is represented by a unique symbol and line color. Each symbol is a single plant measure (panel A ‘Superb’, panel B ‘Stettler’, panel C ‘AAC Viewfield’).

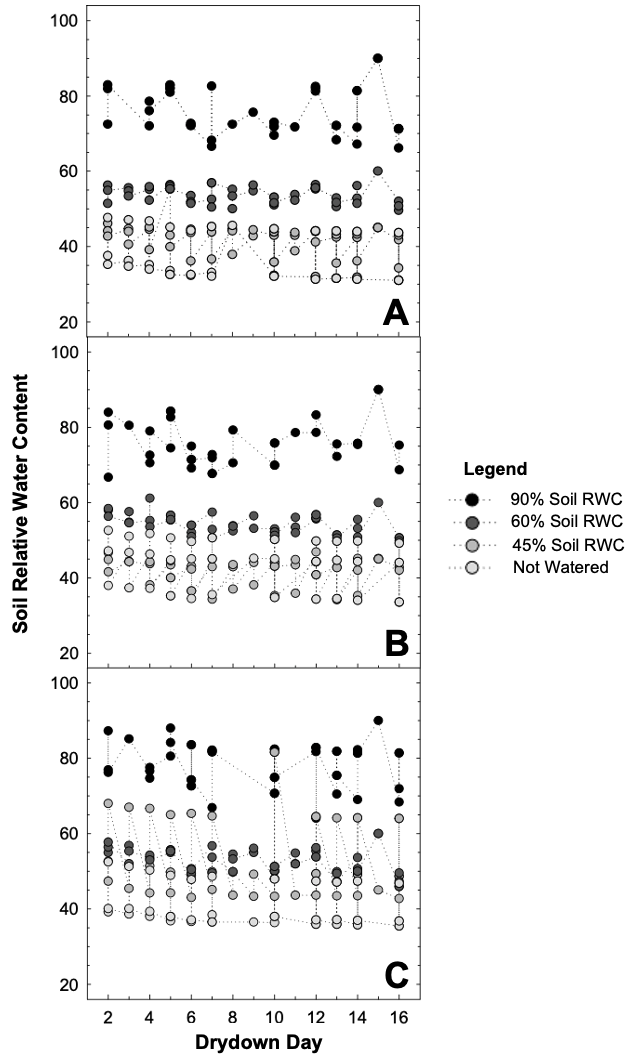
**Supplementary Figure S4.** Temporal changes of soil relative water content of plants measured for experiment 2 (Figures 6 and 7). Each treatment is represented by different symbols and each symbol is the average value of the plants in the treatment (panel A ‘Superb’, panel B ‘Stettler’, panel C ‘AAC Viewfield’). The 90%, 60%, 45% soil relative water content, and not watered treatments are represented by black, dark grey, light grey and white symbols respectively.

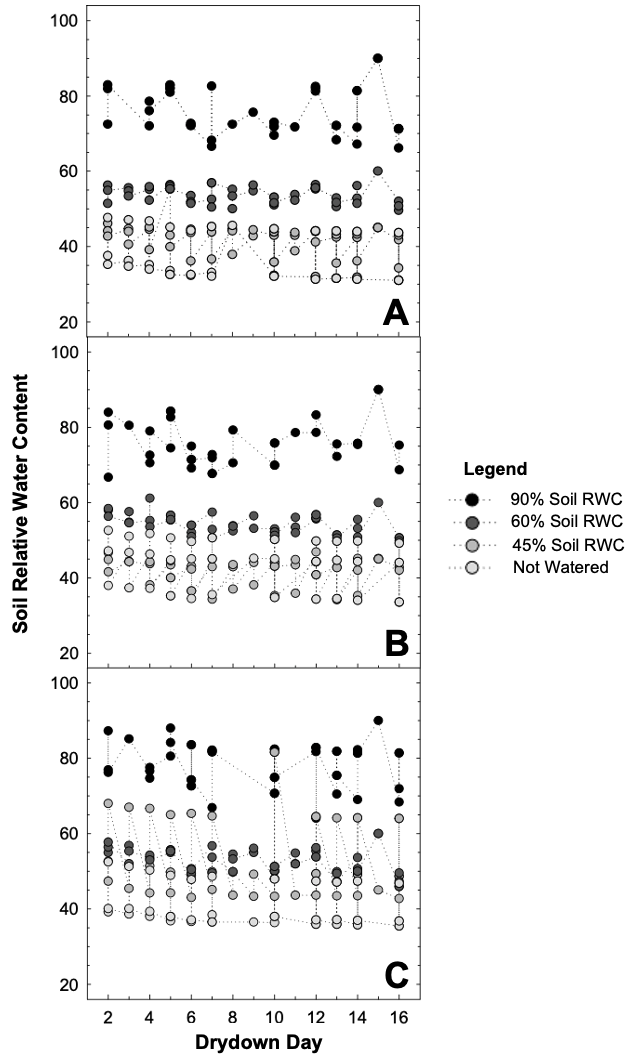

**Supplementary Figure S5.** Representative image of wheat plants growing in large cylindrical plastic pots inside the greenhouse compartment (panel A). Relationship of soil relative water content and gravimetric water content measured from soil cores at top (panel B), middle (panel C) and bottom (panel D) location along pots. Triplets of data points at a given soil relative water content come from the same pot. The top, middle, bottom soil layers are represented by circle, triangle, and square shapes respectively.

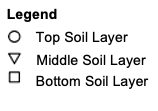

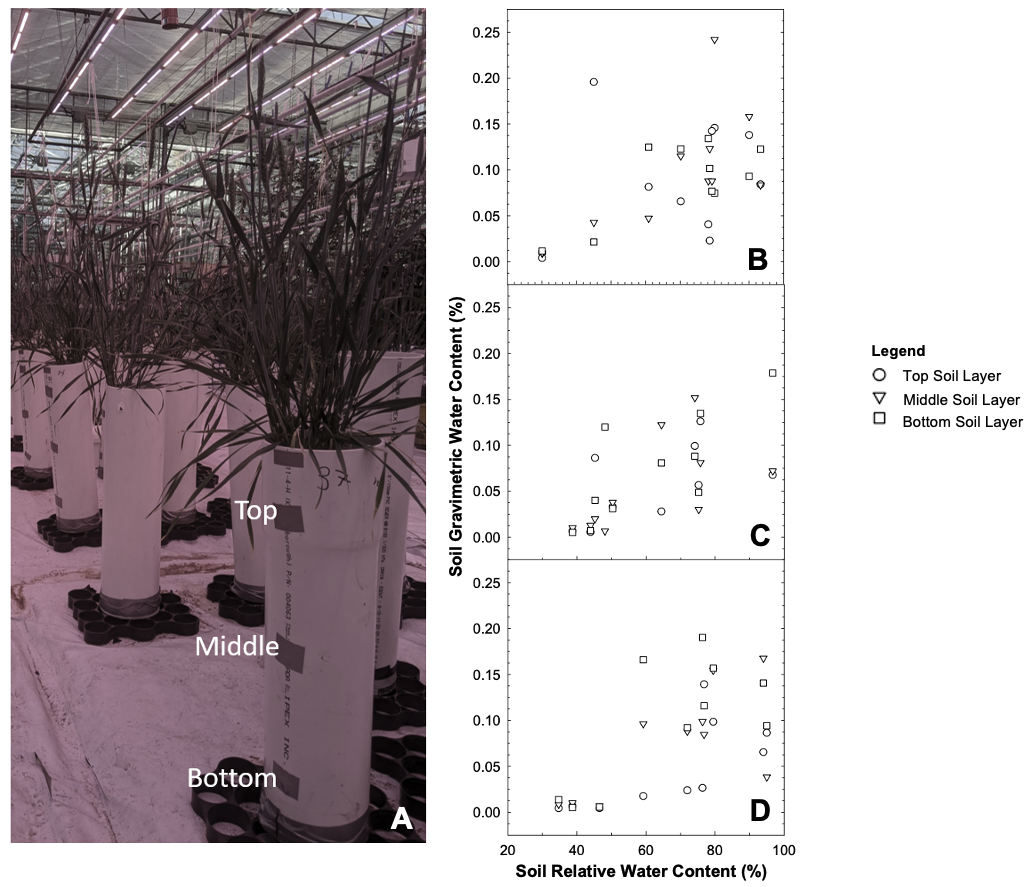

**Supplementary Table S1**. Statistical comparison of soil gravimetric water content (Supplementary Figure S5) using ANOVA analyses (mixed effects linear regression model). Values are the respective F and p-values (Type II) for ‘cultivar’ effect and ‘soil relative water content’ effect. The asterisk (*) indicates *P*<0.05.

| **Parameter** | *Factor* | *F* | *P* |
| --- | --- | --- | --- |
| Soil Gravimetric Water Content | *Cultivar* | 1.1664 | 0.326 |
|  | *Soil Relative Water Content* | 179.1754 | <2e-16* |

**Supplementary Table S2.** Statistical comparison of soil gravimetric water content (Supplementary Figure S5) using ANOVA analyses (mixed effects linear regression model). Values are the respective F and p-values (Type II) for ‘depth’ effect (soil layer) and ‘soil relative water content’ effect. The asterisk (*) indicates *P*<0.05.

| **Parameter** | *Factor* | *F* | *P* |
| --- | --- | --- | --- |
| Soil Gravimetric Water Content | *Depth* | 1.9697 | 0.1425 |
|  | *Soil Relative Water Content* | 163.66 | <2e-16* |

**Supplementary Figure S6.** Overview of experimental procedure for measurement of leaf relative water content, solute potential, and water potential under non-transpiring (left hand panels) and transpiring (right hand panels) conditions.

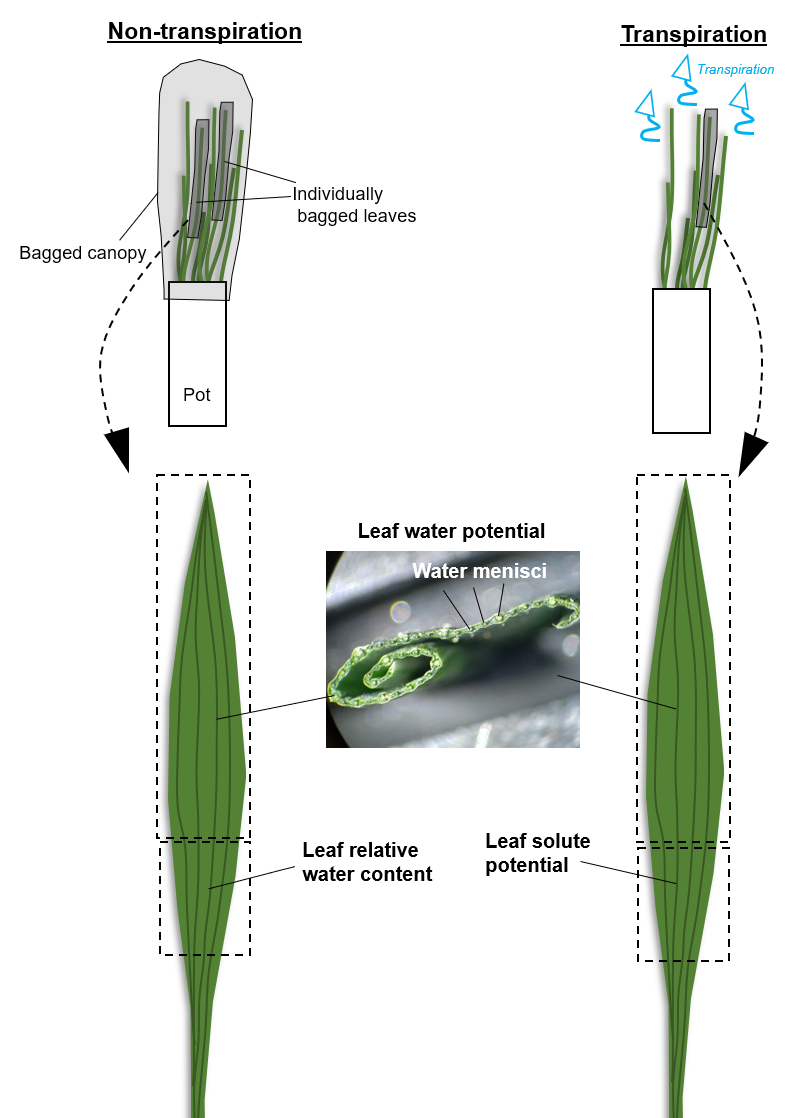

**Supplementary Table S3.** Statistical comparison of a mixed effects linear regression among cultivars using ANOVA analyses (Type III). Analyses of leaf parameters for leaf hydration phase I (i.e., range between 100% to Θ_RWC_). Values are the respective F and p-values (Type III) for ‘cultivar’ effect, ‘soil relative water content’ effect, and interaction effect. The asterisk (*) indicates *P*<0.05.

| **Leaf parameter** | *Factor* | F | *P* |
| --- | --- | --- | --- |
| Leaf relative water content | *Cultivar* | 3.1846 | 0.044208* |
|  | *Soil Relative Water Content* | 9.2351 | 0.003458* |
|  | *Cultivar x Soil Relative Water Content* | 4.5376 | 0.012173* |
| Flag leaf width | *Cultivar* | 13.796 | 2.508e-06* |
|  | *Soil Relative Water Content* | 15.409 | 0.0001213* |
|  | *Cultivar x Soil Relative Water Content* | 15.366 | 6.406e-07* |

**Supplementary Table S4.** Statistical comparison of mixed effects linear regressions among cultivars using a post-hoc Tukey’s analysis (corresponding to Supplementary Table S3). Data shown are for soil relative water content and leaf relative water content in phase I (i.e., range between 100% to Θ_RWC_). Values are predicted p-values for ‘cultivar’ effect. The corresponding mean estimate, standard error of means (SEM), t values and p-values are given for each comparison. The asterisk (*) indicates *P*<0.05.

| **Contrast** | *Mean* | *SEM* | *t* | *P* |
| --- | --- | --- | --- | --- |
| AAC Viewfield – Stettler | 1.22 | 1.10 | 1.109 | 0.5097 |
| AAC Viewfield – Superb | 2.90 | 1.08 | 2.680 | 0.0220* |
| Stettler – Superb | 1.69 | 1.06 | 1.591 | 0.2526 |

**Supplementary Table S5.** Statistical comparison of a mixed effects linear regression among cultivars using ANOVA analyses (Type II). Data shown are for soil relative water content corresponding to phase I (i.e., range between 100% to Θ_RWC_). Values are the respective F and p-values (Type II) for ‘cultivar’ effect and ‘soil relative water content’ effect. The asterisk (*) indicates *P*<0.05.

| **Leaf parameter** | *Factor* | F | *P* |
| --- | --- | --- | --- |
| Solute potential | *Cultivar* | 0.3123 | 0.73128 |
|  | *Soil Relative Water Content* | 3.6601 | 0.05978 |
| Leaf water potential (bagged canopy) | *Cultivar* | 7.7725 | 0.0005766* |
|  | *Soil Relative Water Content* | 0.2940 | 0.5888817 |
| Leaf water potential (transpiring canopy) | *Cultivar* | 3.8392 | 0.02329* |
|  | *Soil Relative Water Content* | 22.9263 | 1.003e-05* |

**Supplementary Table S6.** Statistical comparison of mixed effects linear regressions among cultivars using a post-hoc Tukey’s analysis (corresponding to Supplementary Table S5). Data shown are for soil relative water content corresponding to non-transpiring water potential data in phase I (i.e., range between 100% to Θ_RWC_). Values are predicted p-values for ‘cultivar’ effect and ‘individual’ effect. The corresponding mean estimate, standard error of means (SEM), t values and p-values are given for each comparison. The asterisk (*) indicates *P*<0.05.

| **Contrast** | *Mean* | *SEM* | *t* | *P* |
| --- | --- | --- | --- | --- |
| AAC Viewfield – Stettler | 0.280 | 0.0721 | 3.891 | 0.0004* |
| AAC Viewfield – Superb | 0.142 | 0.0714 | 1.987 | 0.1182 |
| Stettler – Superb | -0.138 | 0.0698 | -1.983 | 0.1193 |

**Supplementary Table S7.** Statistical comparison of mixed effects linear regressions among cultivars using a post-hoc Tukey’s analysis (corresponding to Supplementary Table S5). Data shown are for soil relative water content corresponding to transpiring water potential data in phase I (i.e., range between 100% to Θ_RWC_). Values are predicted p-values for ‘cultivar’ effect and ‘individual’ effect. The corresponding mean estimate, standard error of means (SEM), t values and p-values are given for each comparison. The asterisk (*) indicates *P*<0.05.

| **Contrast** | *Mean* | *SEM* | *t* | *P* |
| --- | --- | --- | --- | --- |
| AAC Viewfield – Stettler | 0.1161 | 0.0561 | 2.069 | 0.0993 |
| AAC Viewfield – Superb | 0.1331 | 0.0559 | 2.380 | 0.0479* |
| Stettler – Superb | 0.0169 | 0.0546 | 0.310 | 0.9483 |

**Supplementary Table S8.** Statistical comparison of mixed effects linear regressions among cultivars using a post-hoc Tukey’s analysis (corresponding to Supplementary Table S3). Data shown are for soil relative water content corresponding to flag leaf width data in phase I (i.e., range between 100% to Θ_RWC_). Values are predicted p-values for ‘cultivar’ effect. The corresponding mean estimate, standard error of means (SEM), t values and p-values are given for each comparison. The asterisk (*) indicates *P*<0.05.

| **Contrast** | *Mean* | *SEM* | *t* | *P* |
| --- | --- | --- | --- | --- |
| AAC Viewfield – Stettler | -0.371 | 0.503 | -0.738 | 0.7410 |
| AAC Viewfield – Superb | -1.378 | 0.477 | -2.887 | 0.0120* |
| Stettler – Superb | -1.007 | 0.497 | -2.026 | 0.1088 |

**Supplementary Table S9.** Statistical comparison of thousand kernel weight among cultivars using ANOVA analyses (Type III). Values are the respective F and p-values (Type III) for ‘cultivar’ effect, ‘treatment’ effect, and interaction effects. The asterisk (*) indicates *P*<0.05.

| **Factor** | *F* | *P* |
| --- | --- | --- |
| *Cultivar* | 3.815 | 0.0364* |
| *Treatment* | 14.746 | 1.19e-05* |
| *Cultivar x Treatment* | 1.916 | 0.1192 |

**Supplementary Table S10.** Statistical comparison of thousand kernel weight among treatments using a post-hoc Tukey’s analysis. Values are predicted p-values for ‘cultivar’ effect and ‘treatment’ effect. The corresponding p-values are given for each comparison. The asterisk (*) indicates *P*<0.05.

| **Contrast** | *P* |
| --- | --- |
| 60%– 45% | 0.0325110* |
| 90% – 45% | 0.4149502 |
| 30% – 45% | 0.0130782* |
| 90% – 60% | 0.5182762 |
| 30% – 60% | 0.0000090* |
| 30% – 90% | 0.0002775* |

**Supplementary Table S11.** Statistical comparison of thousand kernel weight among cultivars using a post-hoc Tukey’s analysis. Values are predicted p-values for ‘cultivar’ effect and ‘treatment’ effect. The corresponding p-values are given for each comparison. The asterisk (*) indicates *P*<0.05.

| **Contrast** | *P* |
| --- | --- |
| AAC Viewfield – Stettler | 0.8472570 |
| AAC Viewfield – Superb | 0.0385183* |
| Stettler – Superb | 0.1177149 |

**Supplementary Figure S7.** Representative pictures of extracted root systems from three wheat cultivars (panel A ‘Superb’, panel B ‘Stettler’, panel C ‘AAC Viewfield’). The maximum length of the root system in pictures.

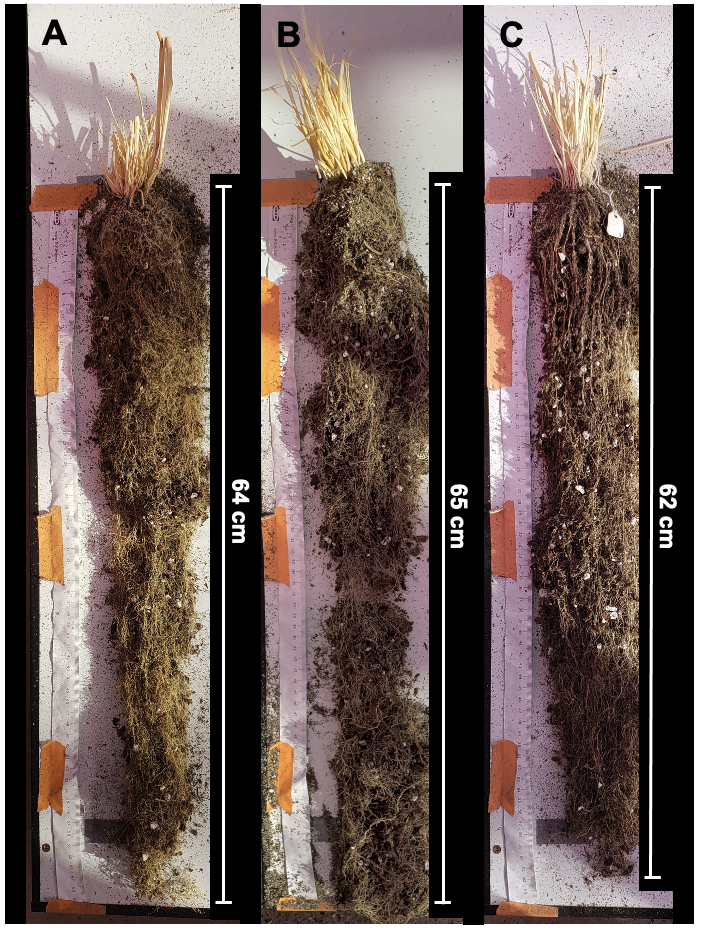

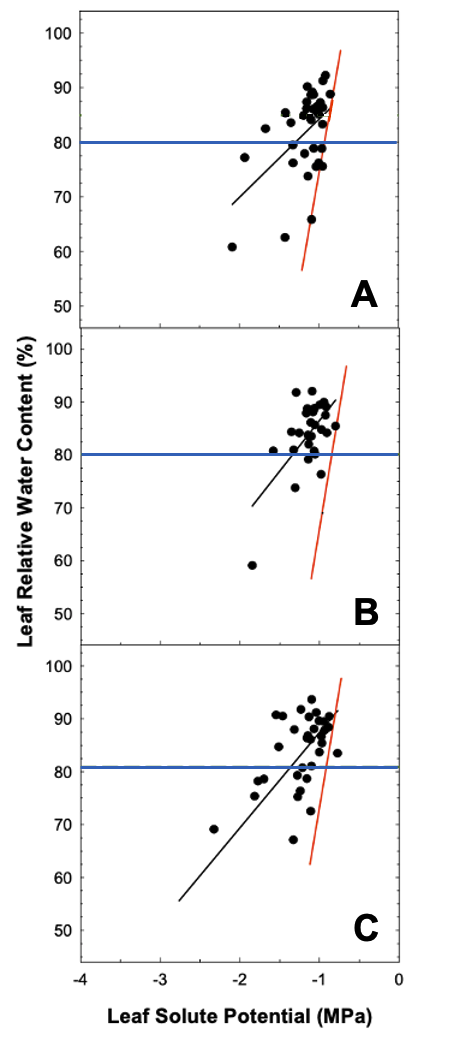
**Supplementary Figure S8.** Relationship of leaf solute potential and leaf relative water content (panel A ‘Superb’, panel B ‘Stettler’, panel C ‘AAC Viewfield’). The dark circular points represent the data gathered during the drydown. The red line represents a model which calculated leaf relative water content for each leaf solute potential point using the method by Babu et al. (1999). Data between 60% and 100% soil relative water conyrny was used to create the linear model of leaf Ψ _S_. A superimposed blue line represents 80% leaf relative water content where the difference between the gathered data and the model was determined. Each symbol represents a single measurement of an individual plant.
